# Apolipoprotein-L1 G1 variant contributes to hydrocephalus but not to atherosclerosis in apolipoprotein-E knock-out mice

**DOI:** 10.1101/2024.12.28.630625

**Authors:** Teruhiko Yoshida, Zhi-Hong Yang, Shinji Ashida, Zu Xi Yu, Shashi Shrivastav, Krishna Vamsi Rojulpote, Piroz Bahar, David Nguyen, Danielle A. Springer, Jeeva Munasinghe, Matthew F. Starost, Victoria J. Hoffmann, Avi Z. Rosenberg, Bibi Bielekova, Han Wen, Alan T. Remaley, Jeffrey B. Kopp

## Abstract

**Introduction:** In USA, six million individuals with Sub-Saharan ancestry carry two *APOL1* high-risk variants, which increase the risk for kidney diseases. Whether APOL1 high-risk variants are independent risk factors for cardiovascular diseases is unclear and requires further investigation.

**Methods:** We characterized a mouse model to investigate the role of APOL1 in dyslipidemia and cardiovascular diseases. Transgenic mice carrying APOL1 (G0 and G1 variants) on bacterial artificial chromosomes (BAC/APOL1 mice) were crossed with the ApoE knock-out (ApoE-KO) atherosclerosis mouse model. The compound transgenic mice were evaluated for the impact of APOL1 on systemic phenotypes.

**Results:** ApoE-KO mice carrying APOL1-G0 and APOL1-G1 did not show differences in the extent of atherosclerotic lesions or aortic calcification, as evaluated by Sudan IV staining and radiographic examination, respectively. However, ∼20% of ApoE-KO; BAC/APOL1-G1 mice developed hydrocephalus and required euthanasia. The hydrocephalus was communicating and likely was due to excess cerebrospinal fluid produced by the choroid plexus, where epithelial cells expressed APOL1. Single-nuclear RNA-seq of choroid plexus identified solute transporter upregulation and mTORC2 pathway activation in APOL1-G1-expressing epithelial cells. Further, in the All of Us cohort, we found higher hydrocephalus prevalence among individuals with the *APOL1-G1* variant in both recessive and dominant models, supporting the mouse findings.

**Conclusion:** While APOL1-G1 expression in ApoE-KO mice did not worsen cardiovascular disease phenotypes, we uncovered hydrocephalus as a novel APOL1 risk allele-mediated phenotype. These findings extend the spectrum of APOL1-associated pathologies.

## Introduction

Two *APOL1* coding variants, termed G1 and G2, explain a substantial proportion of the high prevalence of kidney disease in individuals with sub-Saharan African ancestry.^1–3^ Approximately 13% of African Americans carry two risk variants and have greatly increased risk for chronic kidney disease (CKD).^4^ The utility of APOL1 genotype testing is under investigation in various clinical settings.^5,6^ Further, APOL1 inhibition strategies are being tested in clinical trials of APOL1-mediated kidney diseases.^7–9^

APOL1 is expressed in multiple tissues in addition to kidney, including liver^10^, and vasculature.^11,12^ APOL1 also circulates in a subpopulation of high-density lipoprotein particles. Carriage of two APOL1 risk variants may also be associated with increased risk for cardiovascular diseases (CVD). Studies investigating a possible relationship between the APOL1 risk variants and CVD have reported conflicting results^13^, with some reporting an association^14–18^ and others finding no association.^19,20^ In an autopsy cohort, APOL1 risk variants were independently associated with an increased risk of thrombotic coronary death due to plaque rupture.^21^ However, CKD promotes CVD, which makes it more challenging to establish an independent role for APOL1 in CVD. Indeed, in some reports, an independent association between the APOL1 risk variants and CVD risk is lost when modeled together with CKD.^14,15^ Other studies and meta-analyses found no association between the APOL1 risk variants and the risk of CVD.^19,20,22–25^ Thus, it remains unclear whether APOL1 genotype directly modifies the risk of CVD, independent of its renal effects.

To investigate whether APOL1 plays independent pathogenic role in CVD, we studied a new mouse model: we cross-bred apolipoprotein-E (ApoE) knock-out mice, a well-established atherosclerosis model, with BAC/APOL1 mice, carrying human genomic constructs from bacterial artificial chromosomes (BAC) including APOL1-G0 or G1 alleles. While an APOL1-mediated augmentation of a CVD phenotype was not observed, we unexpectedly discovered a hydrocephalus phenotype in ApoE-KO; BAC/APOL1-G1 mice.

Hydrocephalus imposes a heavy disease burden in sub-Saharan Africa, with ∼200,000 estimated new cases annually among a population of 1.2 billion; this is also a region where APOL1 high-risk variants are common.^26^ Further, African Americans are overrepresented in hydrocephalus-related admissions in the US.^27^ The pathologic mechanisms responsible for hydrocephalus are still incompletely defined, but include alterations in cerebrospinal fluid flow dynamics.^28^ Here we demonstrate a novel role for the APOL1-G1 variant in the pathophysiology of hydrocephalus in transgenic mice, and this genetic association is supported by clinical data.

## Methods

### Mice

Mouse experiments were conducted in accordance with the National Institutes of Health Guide for the Care and Use of Laboratory Animals and were approved in advance by the NIDDK Animal Care and Use Committee (Animal study proposal, K097-KDB-17 & K096-KDB-20).

We studied human *APOL1* gene-locus transgenic mice that contained a bacterial artificial chromosome (BAC) carrying the genomic sequence for APOL1-G0 or G1 (Merck & Co., Inc., Rahway, NJ, USA).^29–31^ These mice have two independent locations of the insertions of transgenes. We purchased ApoE-KO (B6.129P2-*Apoe^tm1Unc^*/J) mice from the Jackson Laboratory, Bar Harbor, ME.^32,33^

Heterozygous BAC/APOL1-G0 or G1 mice were backcrossed to ApoE-KO mice for at least three generations. ApoE-KO, ApoE-KO; BAC/APOL1-G0, and ApoE-KO; BAC/APOL1-G1 mice were identified by genotyping (Transnetyx, Cordova, TN), using targeting probes for ApoE-WT, ApoE-KO, APOL1 (rs60910145 for G0, rs73885319 for G1). Since BAC/APOL1-G0 and BAC/APOL1-G1 mice are heterozygous, we used compound transgenic mice with three genotypes for experiments and with the same background: ApoE-KO, ApoE-KO; BAC/APOL1-G0, and ApoE-KO; BAC/APOL1-G1. Mice were monitored and harvested when reaching an endpoint, defined as a dome-shaped head or alternatively at 9 months of age.

We studied both male and female mice, except for some particular assays, as described below. *APOL1*-transgenic mice and wild-type mice were matched for sex and age. Mice were housed in cages on a 12-hour light/dark cycle, with controlled temperature and humidity. Food and water were provided *ad libitum*. Sample sizes for experiments were determined without formal power calculations.

### Mouse chemistry measurements

Plasma samples were used to measure creatinine, glucose, albumin, and cholesterol profiles, consisting of total cholesterol, triglycerides, high-density lipoprotein (HDL) and low-density lipoprotein (LDL) (VRL Maryland, LLC, Gaithersburg, MD). In addition, plasma lipoprotein cholesterol profiles were analyzed using fast-performance liquid chromatography (FPLC). Pooled plasma (150 µl) from each genotype was injected onto Superose 6 10/300 GL column on an AKTA purifier (AKTA Pure, GE Healthcare, Chicago, IL, USA). Lipoproteins were eluted with buffer, 10 mM Tris, 150 mM NaCl, 0.5 mM EDTA, 0.01% sodium azide, pH 7.4, and 0.5 mL fractions were collected at a flow rate of 0.5 mL/min. Total cholesterol in the FPLC fractions was determined using a cholesterol E kit (NC9138103, Fujifilm Wako Chemicals, Richmond, VA) according to the manufacturer’s instructions.

### Echocardiogram of mouse heart

ApoE-KO (n=5), ApoE-KO; BAC/APOL1-G0 (n=4), and ApoE-KO; BAC/APOL1-G1 (n=4) mice at six months of age were measured. Mice were anesthetized with 1-2% isoflurane during examinations and placed in the supine position over a heated platform, with electrocardiography leads and a rectal temperature probe. Heart images were acquired using the Vevo2100 ultrasound system (VisualSonics, Toronto) with a 30 MHz ultrasound probe (VisualSonics; MS-400 transducer). Measurements were made from standard 2D and M-mode images from the parasternal long axis and midpapillary short axis views of the left ventricle.

### MRI of mouse head

MRI experiments were performed on a 9.4 T 30 cm horizontal scanner (Bruker, Billerica, MA). Mice, anesthetized with an 2% isoflurane/ air mixture, placed on a stereotaxic holder and anesthesia was continued through a nose cone. Core body temperature was maintained at 37°C using circulating water heating pad and was monitored by a rectal temperature probe. A pressure transducer (SA instruments, Stony Brook, NY) was positioned to monitor respiration.

The mouse head was positioned within a 2×2, four-channel receiver array and a 86 mm transmission radio frequency (RF) coil ensemble (Bruker) and centered in the MRI scanner. The RF coil performance and the magnetic field homogeneity, experienced within the brain region, were optimized. A pilot scan of the brain was acquired in three orthogonal planes. Subsequently, T2-weighted 2-dimensional scans (repetition time = 2500 ms, effective echo time = 46 ms, Echo Train length =16, in-plane resolution = 100 mm, 1 mm slices) encompassing the whole brain, were acquired.

### Aorta calcification quantification

The mouse aorta samples underwent X-ray imaging at 10 micrometer resolution on a custom benchtop X-ray microscope to quantify calcification in each sample.^34^ The aorta samples were placed in a flat tray and fully immersed in distilled water during imaging, which allowed a modified single-energy X-ray absorptiometry method to obtain a two-dimensional density distribution of calcium hydroxyapatite, expressed in units of µg/mm^2^.^35,36^ This distribution was then integrated over the field of view to provide a measure of the total amount of calcification in each sample, expressed in units of µg.

### Aorta atherosclerosis characterization

The whole aorta, including the aortic arch, and ascending, descending, thoracic, and abdominal portions, was dissected from the the aortic root to the femoral artery bifurcation. The freshly isolated aorta was stained with 0.5% Sudan IV solution in acetone and 70% ethanol (1:1; v/v) following a brief fixation in 10% neutral buffered formalin, and excess stain was removed using 70% ethanol. After all perivascular adipose tissues were removed, the aorta was cut open longitudinally and mounted onto a glass slide with a cover slide for imaging of the intact aorta with a digital camera connected to a light microscope. Aortic lesion area was analyzed using ImagePro Premier 9.1 (Media Cybernetics, Silver Spring, MD) by an observer masked to sample identity. The extent of aortic atherosclerosis was expressed as percentage of the Sudan IV-positive area relative to the total aortic internal surface area.

### Mouse pathological evaluation

Formalin-fixed, paraffin-embedded (FFPE) mouse brain tissue sections were stained with hematoxylin and eosin for routine histological assessment. FFPE mouse heart tissue was used to characterize aortic sinuses and ventricles. Aortic sinus slides were stained with oil-red O (Histoserv, Germantown, MD) for quantitative assessment of atherosclerosis. Heart ventricular tissue was stained with hematoxylin and eosin and stained with Masson trichrome for quantitative histological assessment including fibrosis.

### Human choroid plexus acquisition

The human choroid plexus was annotated and retrieved by neurologists and a neuropathologist at the Human Brain Collection Core (HBCC) of the National Institute of Mental Health (NIMH), NIH. This was conducted under the Comprehensive Multimodal Analysis of Patients with Neuroimmunological Diseases of the CNS (Clinical protocol 09-I-0032). Upon acquisition, the tissue was immediately preserved in 10% formalin and subsequently prepared for immunohistochemical analysis. Two FFPE sections were obtained for this study; one for hematoxylin and eosin staining, the other for immunohistochemistry probing APOL1 as described in the following section.

### Immunohistochemistry

FFPE tissue sections were deparaffinized and rehydrated. Antigen retrieval was accomplished by heating in a citrate-buffered medium for 15 min in a 99°C hot water bath. Tissues were blocked with 2.5% normal horse serum. Sections were incubated for one hour at room temperature with the following primary antibodies: APOL1 (5.17D12, rabbit monoclonal, 5 μg/ml) that was kindly provided by Genentech (South San Francisco, CA)^37^; or pNKCC1 (ABS1004, Millipore-Sigma; 1:200); or pSPAK (07-2273, Millipore-Sigma; 1:200); or Phospho-SGK1 (Thr256) (44-1260G, Thermo Fisher Scientific, Waltham, MA, 1:50); or pAkt (9271, Cell Signaling, 1:50). For chromogenic detection, sections were processed following protocols for ImmPRESS HRP Horse Anti-rabbit IgG Polymer Kit, Peroxidase and ImmPACT DAB EqV Peroxidase (HRP) Substrate (MP-7401 and SK-4103, Vector Laboratories, Burlingame, CA), and counter-stained with hematoxylin. For fluorescence detection, sections were incubated with Goat anti-Rabbit IgG (H+L) Highly Cross-Adsorbed Secondary Antibody, Alexa Fluor Plus 488 (A32731TR, Thermo Fisher Scientific) and mounted with Antifade Mounting Medium with DAPI (Vector Laboratories).

### Human choroid plexus epithelial cell culture

Human choroid plexus epithelial cells were purchased (1310, ScienCell, Carlsbad, CA). Cells were grown in epithelial cell medium (4101, ScienCell) and on 2 μg/cm^2^ poly-L-lysine coated plates at 37°C, 95% air and 5% CO_2_. Cells were transfected with pCMV-Tet3G plasmid (631335, Takara Bio USA, San Jose, CA) and either of TRE-Empty, TRE-APOL1-G0, TRE-APOL1-G1, or TRE-APOL1-G2 plasmid using Lipofectamine 3000 (Thermo Fisher Scientific). Cells were seeded to 12-well plates in 60% confluency, a day prior to transfection. Transfection reagents were prepared with 1 μg/well of plasmids and were added and incubated for 8 hours. 1 μg/ml of doxycycline hydrochloride (D3072, Millipore-Sigma) were added and incubated for 16 hours to induce APOL1 expression. Cell lysates for protein measurements were prepared by radioimmunoprecipitation assay (RIPA) buffer (20-188, Millipore-Sigma) supplemented with protease and phosphatase inhibitor cocktail (78440, Thermo Fisher Scientific).

### Immunoblotting

Lysates were separated by SDS-polyacrylamide gel electrophoresis (SDS-PAGE) (gradient gel 4-12%, MOPS buffer) and the proteins subjected to Western blotting and blocked for one hour in Intercept (TBS) Blocking Buffer (927-60001, LI-COR, Lincoln, NE). Primary antibodies were Phospho-SGK1 (Thr256) (44-1260G, Thermo Fisher Scientific, 1:1000 dilution), SGK1 (ab43606, Abcam, 1:1000 dilution), APOL1 (3.7D6+3.1C1, Genentech, 1:5000 dilution), β-actin (47778, Santa Cruz Biotechnology, Dallas, TX, 1:5000 dilution). Blots were imaged using the Odyssey infrared scanner (LI-COR).

### Collection of mouse cerebrospinal fluid (CSF) and cytokine measurements

Glass capillary tubes with an inner diameter about 0.2 mm were prepared from thin wall glass capillaries (TW100F-4, World Precision Instruments, Sarasota, FL), using puller (PC-100, Narishige, Amityville, NY). After anesthetizing mice with isoflurane, 0.01 ml of meloxicam (2 mg/ml, ZooPharm, Laramie, WY) and 0.5 ml of saline were injected intraperitoneally. Mice were immobilized with stereotactic instruments (David Kopf, Tujunga, CA) including a nosecone and ear bars. Area of cisterna magna was exposed by cutting 1-1.5 cm of the neck skin with scissors. Muscles were separated with blunt scissors until cisterna magna is visualized. CSF was collected through aspirator tube by puncturing the cisterna magna by glass capillary tubes. Collected CSF samples were diluted and sent for cytokine measurements with Mouse Cytokine/Chemokine 32-Plex Discovery Assay Array (MD32, Eve technologies, Calgary, Canada).

### BAC/APOL1-G1 transgene insertion site detection

The transgene insertion site was detected by Genome Walker Universal Kit (#638904, Takara Bio USA, Mountain View, CA) and followed the manufacturer’s protocol. Genomic DNA was extracted from BAC/APOL1-G1 mouse tail samples. DNA was digested by restriction enzyme Dra I and ligated with adaptors. DNA fragments including transgene insertion sites were amplified by two rounds of PCR with specific primer pairs. DNA bands were cut out by agarose/EtBr gel and Sanger sequencing was conducted. The obtained sequences were used for BLAST search^38^ to find the matching sequence in the mouse genome. The transgene insertion site was examined using the NCBI Genome Data Browser.^39^

### Single-nuclear RNA-seq of choroid plexus

Mouse brain choroid plexus tissue samples were dissected under the microscope immediately after CO_2_ euthanasia. These tissues were snap-frozen on dry ice. Nuclei from the choroid plexus tissues from ApoE-KO and ApoE-KO; BAC/APOL1-G1 mice (n=1 each) were prepared as follows. Snap-frozen choroid plexus tissues were lysed in EZlysis buffer (#NUC101-1KT, Sigma-Aldrich, St. Louis, MO) and homogenized 15 times using a loose Dounce homogenizer and 5 times in a tight pestle. After 5 min incubation, the homogenate was passed through a 40 µm filter (43-50040, PluriSelect, El Cajon, CA) and centrifuged at 500 g at 4°C for 5 min. The pellet was washed with EZlysis buffer and again centrifuged at 500g at 4°C for 5 min. The pellet was resuspended with Dulbecco’s phosphate-buffered saline (DPBS) with 1% FBS to make a final nuclear preparation for loading on to a 10x Chromium Chip G (10x Genomics, Pleasanton, CA) formation of gel beads in emulsion (GEM).

Single nuclear isolation, RNA capture, cDNA preparation, and library preparation were performed following the manufacturer’s protocols for Chromium Next GEM Single Cell 3’ Reagent Kit, v3.1 chemistry (10x Genomics). Prepared cDNA libraries were sequenced at the DNA Sequencing and Genomics Core (NHLBI/NIH). Sequencing and mapping statistics of single-nucleus RNA-seq are presented in **Supplemental Table 1**.

### Single-nuclear RNA-seq analysis

Removal of ambient RNA was accomplished using SoupX (version 1.5.2)^40^ following the default protocol using the autoEstCont and adjustCounts functions. Nuclei with the any of the following features were filtered out: detected gene numbers <200 or >4000, total RNA count >15,000, or mitochondrial transcript percentage >20%. After filtering, 20,996 nuclei remained. Doublets were identified and removed by DoubletFinder (version 2.0.3).^41^

After these pre-processing steps, 18,872 nuclei remained. Integration of single-nucleus gene expression data was performed using Seurat (version 4.0.5) and SeuratData (version 0.2.2).^42^ Clustering was performed by constructing a K-nearest neighbor graph by the first 30 principal components and applying the Louvain algorithm with resolution 0.6. Dimensional reduction was performed with UMAP, resulting in 23 clusters were annotated based on the expression of specific markers.^43,44^ After removing clusters including less than 10 nuclei from both genotypes, 16,641 nuclei from 11 clusters were used for downstream analysis. Genes that were differentially expressed among cell types were assessed with the Seurat FindMarkers function using default parameters. Differentially-expressed genes (DEG) were identified for each paired comparison of genotype, using a cut-off of adjusted P <0.05. Pathway analysis was performed using QIAGEN Ingenuity Pathway Analysis (IPA) software.^45^

### Hydrocephalus and atherosclerosis prevalence and APOL1 genotype analysis in the All of Us Cohort

The All of Us^46^ data set (v6) platform was queried to find individuals who self-identified as African-American and had available genomic data, obtained by either genotyping array or whole genome sequencing (N=54536 participants). All subjects gave informed consent to participate in the All of Us study. The prevalence of hydrocephalus was identified by conditions, as defined in the data set by the following concept identifiers: 4256924, 312902, 440700, 438244, 37204822, 4043738, 4105343, 432899, 440385. Concept identifiers information is summarized in **Supplemental Table 2**. The prevalence of atherosclerosis was identified by the following conditions: concept id 44825446, 1569271, 44823124, 35207841, 45567167, 764123, 1569274, 44828992, 1569272, 44825447, 1569146, 317576, 44835931, 45567168, 44830080, 44835932, 1569320, 45581810, and 1569273. Concept identifiers information is summarized in **Supplemental Table 3**.

APOL1 gene haplotypes are composed of three polymorphisms: G1 (rs73885319 and rs6091945, both missense variants); G2 (rs71785313, a 6 base-pair deletion); and G0, the ancestral haplotypes which does not carry the G1 or G2 variants. APOL1 genotypes were defined by the presence or absence of variant alleles (rs73885319 (G1) and rs71785313 (G2) using Hail software (version 0.2.107).

The associations between *APOL1* genotype and the prevalence of hydrocephalus and atherosclerosis were analyzed. Univariate analyses to calculate odds ratios, p-values, and 95% CI were performed using the Fisher’s exact test (for APOL1 genotype model and sex) or t-test (for age and eGFR, continuous variables), implemented within the R stats (version 3.6.2) package. Multi-variate logistic regression analyses were performed with adjustment of age, sex, and estimated glomerular filtration rate (eGFR). P < 0.05 was considered statistically significant. Code is available, upon request, to All of Us users with access to the controlled tier data set.

## Results

### Communicating hydrocephalus development in ApoE-KO/APOL1-G1 mice

We generated and characterized ApoE-KO, ApoE-KO; BAC/APOL1-G0, and ApoE-KO; BAC/APOL1-G1 mice to evaluate the effect of *APOL1* risk genotype superimposed on the phenotype of ApoE-KO mice. ApoE-KO; BAC/APOL1-G1 had shorter survival compared with mice of other genotypes (P=0.018) (**Figure 1A**). Unexpectedly, 21% (3/14) of ApoE-KO/APOL1-G1 mice developed hydrocephalus. This manifested grossly as enlarged “domed” head. MRI and pathology of the brain showed patent fourth ventricles, consistent with communicating hydrocephalus (**Figure 1B**, **1C**).

**Figure 1.**
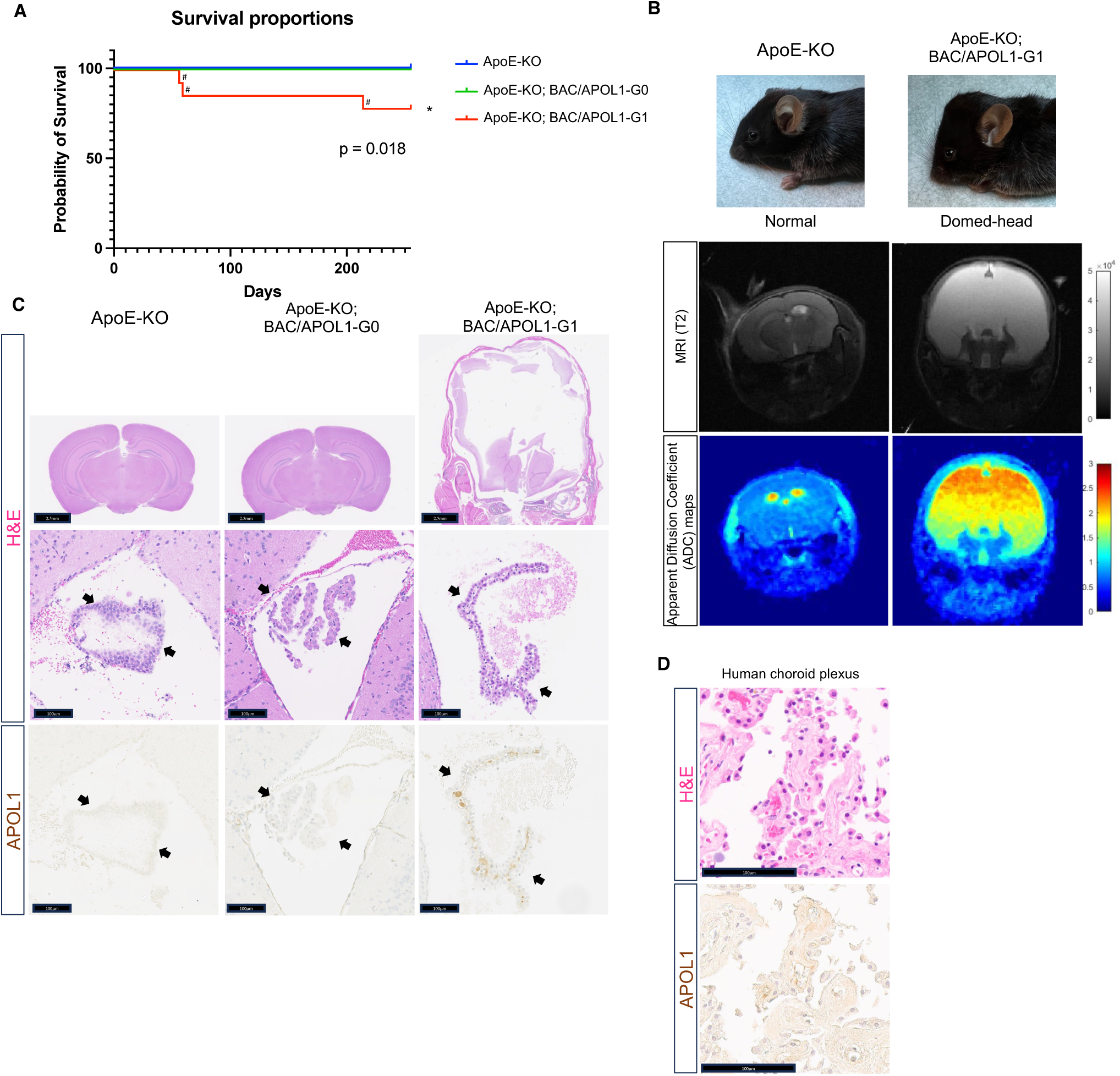
APOL1-G1 variant contributes to hydrocephalus. **(A**) Kaplan-Meier survival curve [ApoE-KO (n=23), ApoE-KO; BAC/APOL1-G0 (n=12), ApoE-KO; BAC/APOL1-G1 (n=14)] showed early death of ApoE-KO; BAC/APOL1-G1 mice (log rank: p = 0.018). ^#^mark indicates mice requiring euthanasia due to hydrocephalus development. (**B**) Characteristic domed-head appearance and magnetic resonance imaging (MRI) T2 weighted images show hydrocephalus in ApoE-KO; BAC/APOL1-G1 mice. (**C**) Representative images of hematoxylin & eosin staining and APOL1 immunohistochemistry, showing APOL1 expression in the choroid plexus epithelium in ApoE-KO; BAC/APOL1 mice. (**D**) Representative images of hematoxylin & eosin staining and APOL1 immunohistochemistry, showing APOL1 expression in human choroid plexus epithelium. KO, knock-out; Scale bars are included in figures. Arrowheads point to choroid plexus epithelium.

One cause of hydrocephalus is the overproduction of cerebrospinal fluid by the choroid plexus. When we characterized the expression of APOL1 by immunohistochemistry, epithelial cells in the choroid plexus expressed APOL1. The human protein atlas^47^ also reports that human choroid plexus expresses *APOL1* mRNA, and we found that it also contains APOL1 protein by immunostaining (**Figure 1D**).

### Upregulation of solute transporters in choroid plexus epithelial cells expressing APOL1-G1

We hypothesized that APOL1-G1 expressed in the choroid plexus tissue caused overproduction of cerebrospinal fluid, resulting in communicating hydrocephalus. To profile transcriptomic signatures of the choroid plexus at single cell resolution, we conducted single-nuclear RNA-seq of choroid plexus tissues (**Figure 2A**). ApoE-KO; BAC/APOL1-G1 mouse with hydrocephalus (n=1) and ApoE-KO mouse without hydrocephalus (n=1) in the littermates were used for the analysis. We analyzed 16,641 nuclei in 11 clusters, as identifed by unbiased clustering (**Figure 2B**). Each cluster was annotated using marker genes as reported previously.^43,44^ Marker genes included *Folr1* and *Prlr* for choroid plexus epithelial cells, *S100b* and *Gfap* for astrocytes, *Rbfox3* and *Map2* for neurons, *Mog* for oligodendrocytes, *Slc1a3* for multiple glial cells (astrocytes, microglia, oligodendrocytes and oligodendrocyte precursors), and *Ptprc* for border-associated macrophages. Choroid plexus epithelial cells showed expression of APOL1 (**Figure 2B**).

**Figure 2.**
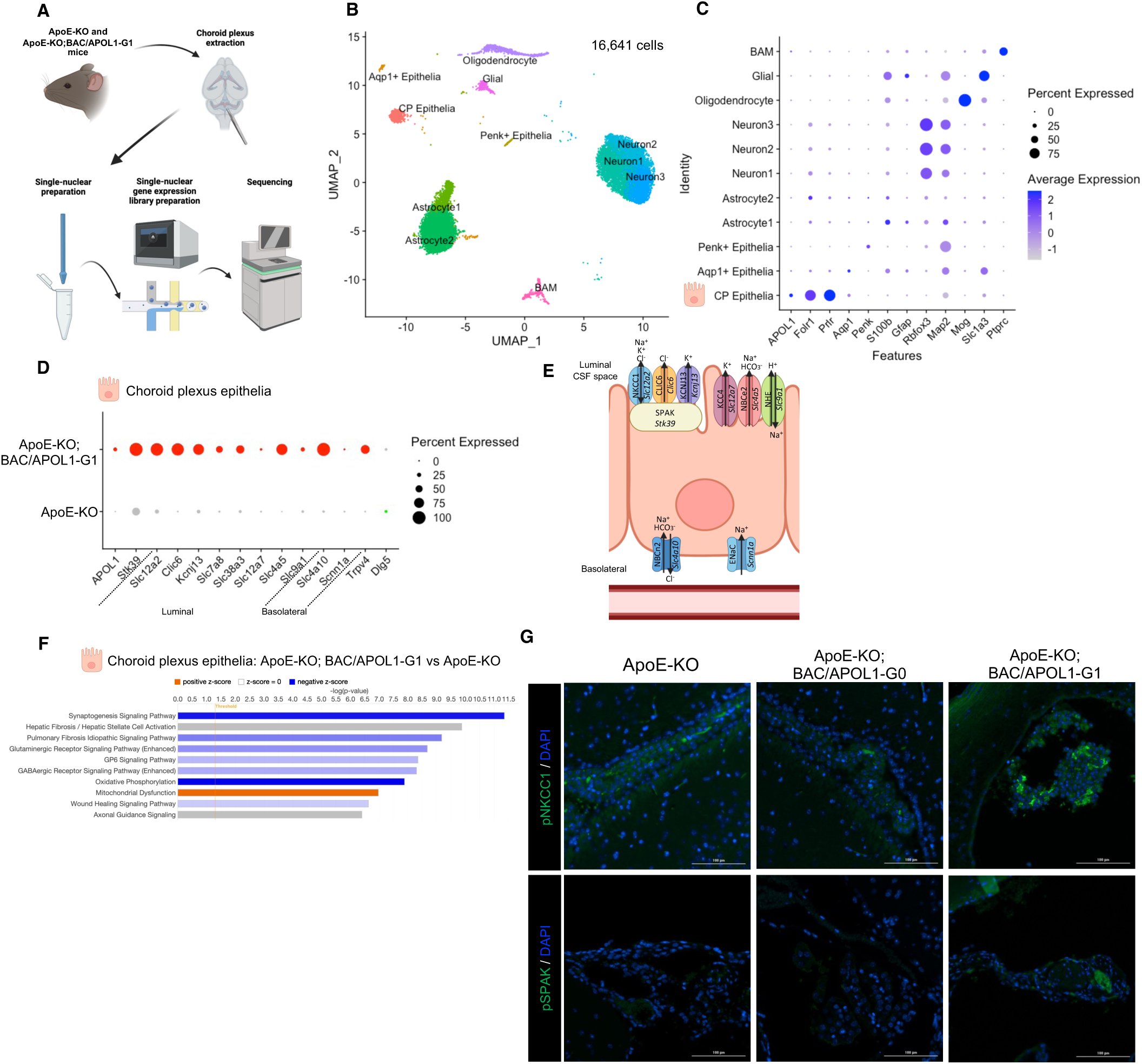
Single-nucleus RNA-seq analysis of choroid plexus showed upregulation of transporters in choroid plexus epithelial cells. **(A**) Schematic of single-nucleus RNA-seq experiments using mouse choroid plexus tissue. (**B**) UMAP plot of single-nuclear RNA-seq data, showing 16,441 cells, distributed among 11 distinct cell clusters. (**C**) Dot plot shows marker genes characteristic of each of 11 clusters. (**D**) Dot plot shows the upregulated expression of solute transporters in choroid plexus epithelium from ApoE-KO; BAC/APOL1-G1 mice, compared to APOE-KO mice. (**E**) Schematic of representative upregulated genes in ApoE-KO; BAC/APOL1-G1 mice shows localization in choroid plexus epithelial cells. (**F**) Pathway analysis of differentially-expressed genes in choroid plexus epithelium, comparing ApoE-KO; BAC/APOL1-G1 mice with APOE-KO mice. With regard to ApoE-KO; BAC/APOL1-G1 mice, orange denotes activated pathways, blue denotes deactivated pathways, and gray denotes unavailable pathway activation score. (**G**) Immunofluorescent images detected phospho-NKCC1 (Na-K-Cl co-transporter) and phospho-SPAK (STE20/SPS1-related proline/alanine-rich kinase), showing higher expression in ApoE-KO; BAC/APOL1-G1 mouse choroid plexus. CP, choroid plexus; BAM, border-associated macrophages

When evaluated DEG in choroid plexus epithelial cells, genes encoding solute transporters reported to be upregulated in the hydrocephalus model^48^, forming complex together with SPAK (*Stk39*), including NKCC1 (*Slc12a2*), CLIC6 (*Clic6*) and KCNJ13 (*Kcnj13*) were more abundant in ApoE-KO; BAC/APOL1-G1 mice (**Figure 2D**, **2E**). ApoE-KO; BAC/APOL1-G1 mice manifested higher expression of other transporters located on the luminal side of the choroid plexus epithelium, including LAT2 (*Slc7a8*) and SNAT3 (*Slc38a3*)^49^ which transport amino acids, KCC4 (*Slc12a7*), NBCe2 (*Slc4a5*) and NHE (*Slc9a1*) which transport solutes (**Figure 2D**, **2E**). Solute transporters located mainly in basolateral side of epithelium include NBCn2/NCBE (*Slc4a10*)^50^, ENaC (*Scnn1a*)^50^, which were more abundant in ApoE-KO; BAC/APOL1-G1 mice (**Figure 2D**, **2E**).

Further, DLG5 (*Dlg5)* maintains polarity of epithelial cells^51^, and Dlg5-/- mice cause hydrocephalus.^52^ This *Dlg5* was downregulated in ApoE-KO; BAC/APOL1-G1 mice. TRPV4 (*Trpv4*) increases secretion of CSF^53,54^, which was also upregulated in choroid plexus epithelial cells in ApoE-KO; BAC/APOL1-G1 mice (**Figure 2D**).

In APOL1-G1 mice, pathway analysis based on the DEGs in choroid plexus epithelial cells showed downregulation of the oxidative phosphorylation pathway, together with upregulated expression of the mitochondrial dysregulation pathway genes (**Figure 2F**). Mitochondrial dysfunction has been described in APOL1 risk-variant expressing cells.^55–57^ In choroid plexus epithelium of ApoE-KO; BAC/APOL1-G1 mice, immunohistochemistry confirmed higher expression of phospho-NKCC1 and phospho-SPAK compared to wild-type mice, as also reported in other hydrocephalus mice^44^ (**Figure 2G**).

### mTORC2 activation in choroid plexus epithelial cells with APOL1-G1 expression

Upstream analysis based on the DEGs in choroid plexus epithelial cells indicated that RICTOR, a component of mTORC2, was the most activated upstream regulator in ApoE-KO; BAC/APOL1-G1 mice compared with ApoE-KO mice (**Figure 3A**). The mTORC2 pathway and SPAK/NKCC1 pathway are known to regulate transporter expression.^58–61^ Therefore, this ApoE-KO; BAC/APOL1-G1 mice is suggested to have mTORC2 pathway activation resulting in transporters’ upregulation intermediated by SPAK activation. Further, the mTORC2 activation was confirmed by measuring higher level of downstream signaling, such as phospho-SGK1 and phosphor-Akt (**Figure 3B**, 3C). This finding is compatible with previous reports showing mTOR activation in hydrocephalus mouse models.^44^

**Figure 3.**
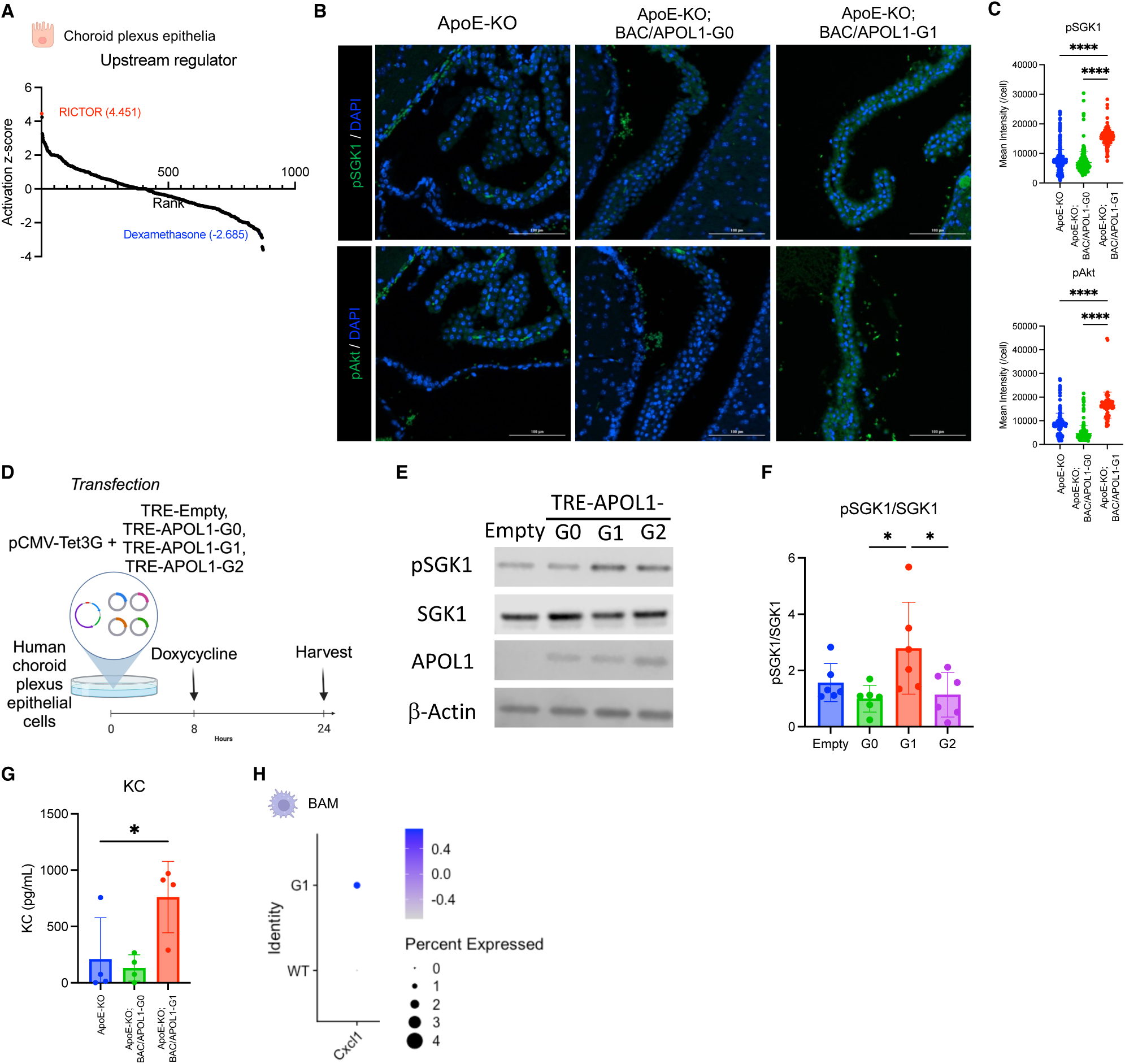
The mTORC2 pathway was upregulated in choroid plexus epithelium of APOL1-G1 mice. **(A**) Waterfall plot shows the results of upstream analysis from differentialy-expressed genes in choroid plexus epithelium, comparing ApoE-KO; BAC/APOL1-G1 mice with APOE-KO mice. RICTOR, a component of mTORC2, was predicted to be the most activated upstream regulator of choroid plexus epithelial cells. (**B**) Immunofluorescent staining showed increased abundance of phospho-SGK1 (top row) and phospho-Akt (bottom row) in ApoE-KO; BAC/APOL1-G1 mouse choroid plexus. **(C**) Measurements of phospho-SGK1 and phospho-Akt immunofluorescence are shown. (**D**) Schematic shows the flow of human choroid plexus epithelial cells experiments. (**E**) Representative immunoblotting images against phospho-SGK1, SGK1, APOL1, and β-actin showed upregulated phosphorylation of SGK1, immediate downstream of mTORC2, in TRE-APOL1-G1 transfected human choroid plexus epithelial cells. (**F**) Quantitative results of immunoblotting (n=4 each) showed increased phospho-SGK1/SGK1 in TRE-APOL1-G1 cells, indicating mTORC2 activation. (**G**) Keratinocyte-derive cytokine (KC) levels in CSF were higher in ApoE-KO; BAC/APOL1-G1 mice compared to other genotypes. (**H**) Dot plot shows higher *Cxcl1* cytokine expression in border-associated macrophages in ApoE-KO; BAC/APOL1-G1 mice compared to APOE-KO mice. Analysis by one-way ANOVA. *, P<0.05; ****, P<0.0001

We used human choroid plexus epithelial cells to investigate whether APOL1 variants induction activates mTORC2. (**Figure 3D**) When APOL1 variants were induced, mTORC2 activation was observed specifically by APOL1-G1 induction. (**Figure 3E**, **3F**)

### Cxcl1/KC (keratinocyte-derived chemokine) as the intermediating cytokine of hydrocephalus in ApoE-knockout/APOL1-G1 mice

We also conducted CSF cytokine measurements from each mouse model (n=4 each) to find markers of hydrocephalus. *Cxcl1*/KC, IL12p70, IL17, *Cxcl10*/IP10 and *Cxcl5*/LIX were upregulated in CSF from ApoE-KO; BAC/APOL1-G1 mice (**Figure 3G, Supplemental Figure S3**). Of these cytokines, *Cxcl1*/KC was reported to be elevated in CSF from idiopathic normal pressure hydrocephalus patients.^62^ Border-associated macrophages from ApoE-KO; BAC/APOL1-G1 mouse showed higher expression of *Cxcl1* compared with ApoE-KO (n=1 mouse each) (**Figure 3H**). Therefore, border-associated macrophages can be the source of KC, augmenting inflammatory responses in the choroid plexus of ApoE-KO; BAC/APOL1-G1 mice.

### Plasma lipids and atherosclerosis in APOL1-G1 mice

Characterization of 9 month-old mice (n=11 (ApoE-KO), n=10 (ApoE-KO; BAC/APOL1-G0 and ApoE-KO; BAC/APOL1-G1)) did not reveal any differences among APOL1 genotypes in body weight (**Supplemental Figure S4**) and serum levels of lipids, including total cholesterol, free cholesterol, HDL cholesterol, LDL cholesterol, triglycerides, and phospholipid. (**Figure 4A-F**). No changes were observed in total cholesterol concentrations in FPLC fractions, including VLDL, LDL, and HDL, among ApoE-KO, ApoE-KO; BAC/APOL1-G0 and ApoE-KO; BAC/APOL1-G1 mice. (**Figure 4G**).

**Figure 4.**
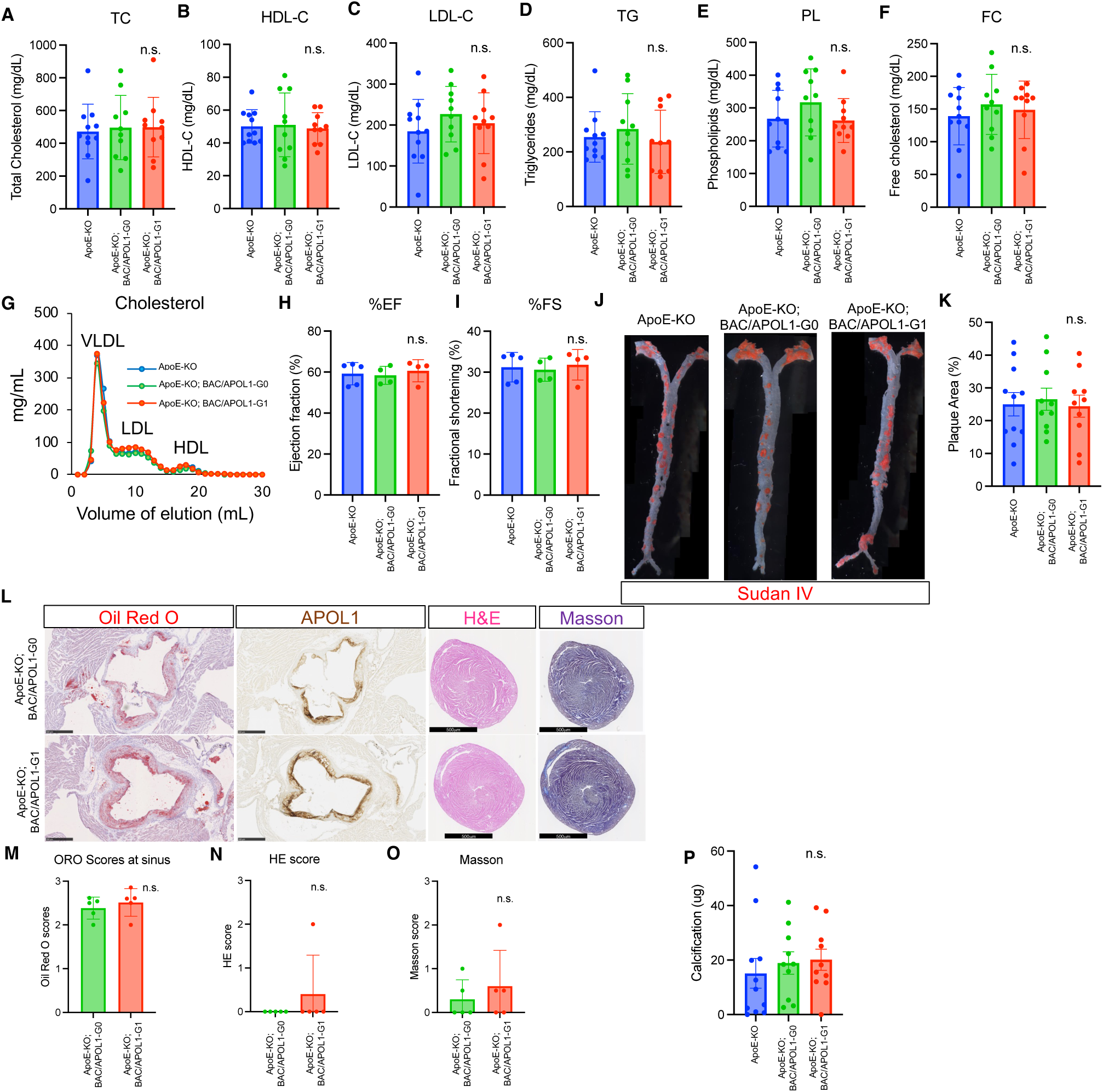
APOL1-G1 expression did not worsen cardiovascular phenotypes in ApoE-KO mice. **(A-F**) Lipid measurements of total cholesterol (TC), high-density lipoprotein cholesterol (HDL-C), low-density lipoprotein cholesterol (LDL-C), triglycerides (TG), phospholipid (PL) and free cholesterol (FC). (**G**) Fast protein liquid chromatograph (FPLC) results of cholesterol in the very-low-density lipoprotein (VLDL), LDL, and HDL fractions. (**H, I**) Ejection fraction and fractional shortening measurements obtained from mouse echocardiography are shown for each mouse genotype. (**J**) Sudan IV images of whole dissected aorta are shown for each mouse genotype. (**K**) Quantitative results of aortic plaque area stained by Sudan IV are shown for each mouse genotype. (**L**) Representative pathology images of oil red O, APOL1 IHC of aorta sinus, and hematoxylin and eosin and Masson staining of cardiac ventricles. Scale bars are 500μm. (**M**) Quantitative scores of oil red O staining of the aortic sinus. (**N, O**) Quantitative scores of H&E and Masson staining of cardiac ventricles showed no differences between groups. (**P**) Quantitative results of aortic calcification measurements showed no differences among groups.

As heart failure promotes atherosclerosis progression in ApoE-KO mice,^63^ echocardiography was performed (n=5 (APOE-KO), n=4 (ApoE-KO; BAC/APOL1-G0 and ApoE-KO; BAC/APOL1-G1)). We did not find any differences in cardiac functional parameters among genotypes (**Figure 4H**, **4I**). With regard to structural parameters, Sudan IV lipid staining of the whole aorta *en face* revealed substantial amounts of plaque (averaging ∼25% of the aorta) in all three mouse groups (**Figure 4J**, **4K**). Oil red O also stained plaque within the aortic sinus, indicating lipid deposition. The extent of atherosclerosis development was the same among genotypes (**Figure 4L**, **4M**). There were no significant cardiac hypertrophy or ventricular fibrosis (**Figure 4L**, **4N**, **4O)**. In addition, similar calcification in aortic plaque was observed in all mouse groups (**Figure 4P**).

### Prevalence of hydrocephalus was higher in individuals with APOL1-G1 genotype in the All of Us cohort

To determine whether hydrocephalus prevalence was higher in individuals with APOL1 high-risk variants, we made use of the All of Us cohort data. Data were analyzed from self-identified African-American individuals with available genotyping array data or whole genome sequencing data (N=54536 participants). The prevalence of hydrocephalus was identified by conditions, as defined in the data set by the concept identifiers summarized in **Supplemental Table 2**. The hydrocephalus prevalence was 0.50% in *APOL1*-G0/G0 individuals (serving as reference), 0.65% in *APOL1*-G0/G1 individuals (odds ratio 1.31 [0.99-1.74], P=0.057), and 0.90% in *APOL1*-G1/G1 individuals (odds ratio 1.85 [1.16-2.86], P=0.0092) (**Supplemental Figure 5A**). For *APOL1*-G1, both the recessive model (odds ratio 1.73 [1.11-2.60], P=0.012) and the dominant model (odds ratio 1.30 [1.03-1.64], P=0.025) showed increased prevalence of hydrocephalus (**Supplemental Figure 5A, 5B**). With the adjustment by age, sex, and eGFR, both the recessive model (odds ratio 4.35 [1.93-8.88], P= 0.00014) and the dominant model (odds ratio 3.86 [2.09-7.50], P= 0.0000028) showed further increased risk of hydrocephalus (**Figure 5A, Supplemental Figure 5C, 5D**). These data indicated that the APOL1-G1 protein acts in both recessive and dominant fashion to promote the development of hydrocephalus in humans, which is consistent with the mouse findings.

**Figure 5.**
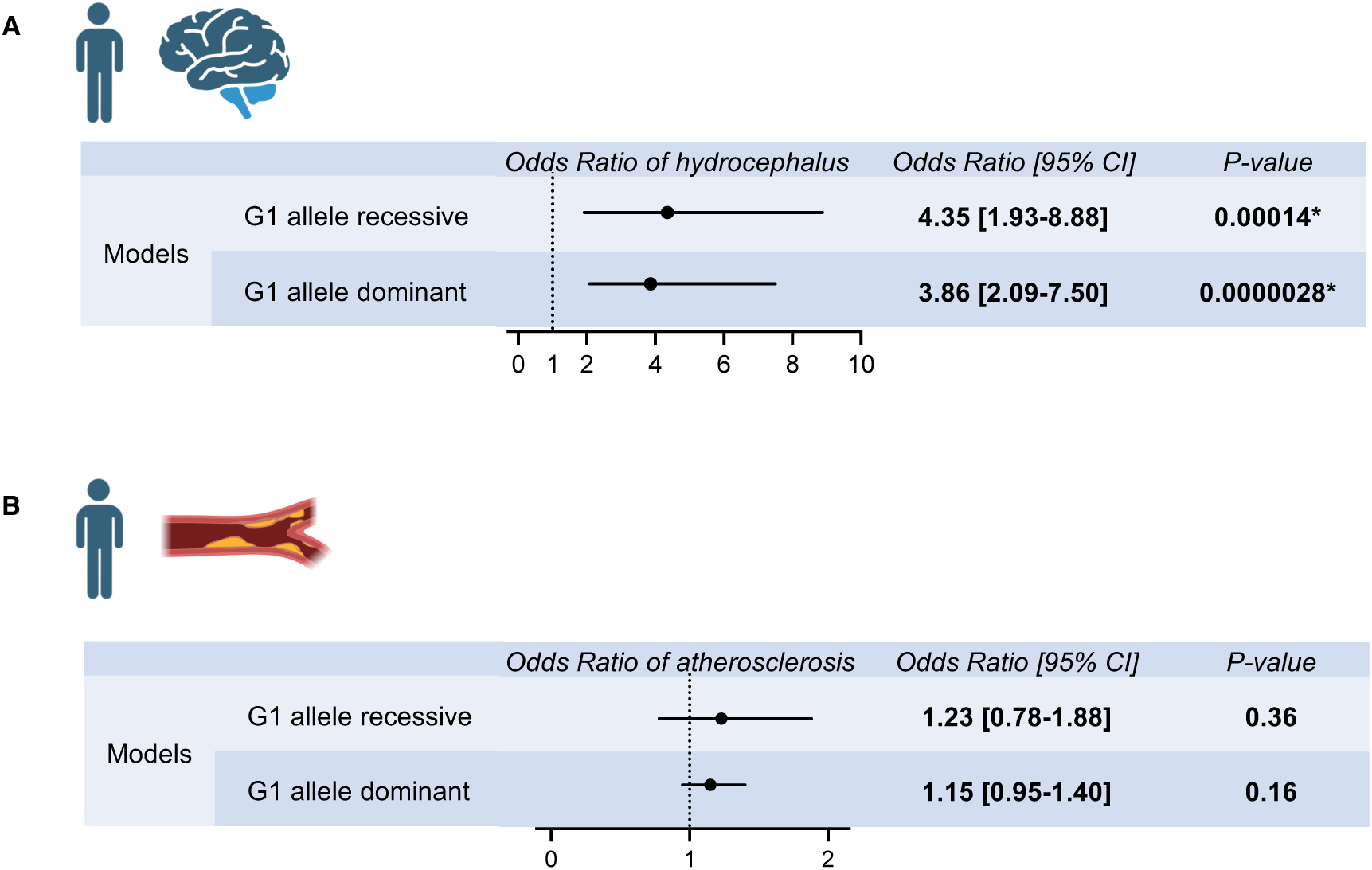
Hydrocephalus and atherosclerosis risks by APOL1-G1 variant in the All of Us population cohort. **(A)** Multi-variate logistic regression analysis results for hydrocephalus odds ratios by APOL1-G1 recessive and dominant models, adjusted for age, sex, and eGFR were shown. APOL1-G1 variant was associated with an increased risk of hydrocephalus. **(B)** Multi-variate logistic regression analysis results for atherosclerosis odds ratios by APOL1-G1 recessive and dominant models, adjusted for age, sex, and eGFR were shown. APOL1-G1 variant was not associated with an increased risk of atherosclerosis.

Further, the prevalence of atherosclerosis was identified by conditions, as defined in the data set by the concept identifiers summarized in **Supplemental Table 3**. The atherosclerosis prevalence was 6.4% in *APOL1*-G0/G0 individuals (serving as reference), 6.1% in *APOL1*-G0/G1 individuals (odds ratio 0.95 [0.87-1.04], P=0.25), and 7.4% in *APOL1*-G1/G1 individuals (odds ratio 1.17 [1.00-1.36], P=0.048) (**Supplemental Figure 6A**). As atherosclerosis was strongly associated with age and eGFR, we conducted the multi-variate analysis adjusting these factors and sex. Although the *APOL1*-G1 recessive model (odds ratio 1.19 [1.02-1.37], P=0.024) showed the increased prevalence of atherosclerosis in univariate analysis (**Supplemental Figure 6A, 6B**), the effect of *APOL1*-G1 genotype on atherosclerosis risk was not observed in recessive model (odds ratio 1.23 [0.78-1.88], P=0.36) by multi-variate analysis with adjustments (**Figure 6, Supplemental Figure 6C, 6D**). With regard to atherosclerosis also, All of Us data for atherosclerotic disease supported the APOL1 mouse model observation.

## Discussion

To investigate the effect of *APOL1* genotype in cardiovascular diseases, we established and characterized a mouse model, by crossing ApoE knock-out mice and BAC/APOL1 mice carrying the *APOL1-G0* or *APOL1-G1* variant controlled by the human APOL1 gene locus promotor. The development of communicating hydrocephalus and resulting early death in ApoE-KO; BAC/APOL1-G1 mice was a novel and unexpected finding. Careful surveillance was needed to detect hydrocephalus prior to death. When hydrocephalus was suspected by a dome-head appearance, the experimental protocol required euthanasia. Necropsy revealed communicating hydrocephalus. Of seven mice sent to necropsy with the dome head phenotype in the entire study, all had hydrocephalus; two mice had brain hemorrhage, and the other two mice had meningitis.

Hydrocephalus is characterized by the expansion of the cerebral ventricles. Congenital hydrocephalus is uncommon but not rare, affecting ∼1 in 1000 live births.^64^ Although classically understood as a disorder of impaired CSF flow, the mechanisms of hydrocephalus are diverse and not fully understood.^65^ Approximately 20% of congenital hydrocephalus cases are explained genetically, but most sporadic cases remain unexplained. Post-infectious hydrocephalus and post-hemorrhagic hydrocephalus mouse models indicate that both lipopolysaccharide and blood breakdown products trigger TLR4-dependent immune responses in the choroid plexus.^44^ There was no modality available to measure absorption of CSF in mice, which limited our research in further understanding of CSF clearance in these mice.

Several pathways may be involved in the development of hydrocephalus in the ApoE-KO; BAC/APOL1-G1 mouse model. The data presented here indicate activation of the mTORC2-SGK1 pathway, leading to upregulated expression of transporters in choroid plexus epithelial cells in ApoE-KO; BAC/APOL1-G1 mice, such as NKCC1 and TRPV4. Human choroid plexus epithelial cell showed mTORC2 activation when APOL1-G1 was induced, which indicated that APOL1-G1 expression in choroid plexus epithelial cell activated mTORC2. It is reasonable to speculate that mTORC2 activates SPAK/OSR1 through phosphorylation as shown in HeLa cells.^66^ ApoE-KO; BAC/APOL1-G1 mouse choroid plexus epithelial cells showed increased abundance and activation of SPAK, together with solute transporters forming complex with. SPAK is a well characterized intermediating factor in regulating NKCC2 (*Slc12a1*) and NCC (*Slc12a3*) expression in renal distal convoluted tubule and thick ascending limb.^67^ Here we show that the expression of APOL1-G1 activates mTORC2-SPAK pathway in choroid plexus epithelial cells, leading to hydrocephalus.

High-risk APOL1 proteins have been implicated in plasma membrane pore formation ^68,69^ and in APOL1-HDL biochemical profile change.^70^ Further research is warranted to understand the role of these and the subcellular localization of APOL1 in choroid plexus epithelial cells. APOL1 protein circulating in CSF^71^ may also play a role in hydrocephalus formation. Untreated infant hydrocephalus in sub-Saharan African countries may result in annual economic losses of more than 50 billion dollars.^26,72^ Although infection is the major cause of infant hydrocephalus,^72^ the role of APOL1 high-risk variants in hydrocephalus needs further investigation. We cannot exclude the remote possibility that the BAC insertion site contributes to some portion of hydrocephalus development in ApoE-KO; BAC/APOL1-G1 mice. However, none of the three neighboring genes (*Hsd17b3, Slc35d2, Zfp367*) within 100 kb of BAC/APOL1-G1 insertion site (mm39, chr13: 64,197,419) were plausibly related to hydrocephalus (**Supplemental Figure 7A**). The expression of those genes was neither abundant nor different between genotypes in choroid plexus single-nucleus RNA-seq data (**Supplemental Figure 7B**). This indicates that a particular insertion site likely does not contribute to hydrocephalus.

All of Us cohort data showed an effect of APOL1-G1 variant on the higher prevalence of hydrocephalus in both recessive and dominant models, which was congruent with the present mouse model. Further validation of this finding in another cohort is warranted. Low penetrance of hydrocephalus in ApoE-KO; BAC/APOL1-G1 mice was not clear, however, unknown environmental factors might have contributed. One possibility is the environmental factors, which induce APOL1 expression. Hydrocephalus is reported in ApoE-KO mice^73^ and 15% of ApoE-KO/LDLR-KO mice^74^. Therefore, the dual transgenic mouse model in this paper likely showed synergistic effect of ApoE-KO and *APOL1* variants. We have not studied APOL1-G2 mice, as BAC/APOL1-G2 mice show lower expression of APOL1 compared with BAC/APOL1-G0 and -G1 mice due to unknown reasons (data not shown).

Cardiovascular disease assessed by atherosclerosis and aortic calcification was not exacerbated by the presence of either APOL1-G0 or APOL1-G1 variants in the ApoE-KO mice. Further, cholesterol profiles and echocardiography did not show differences among *APOL1* genotypes. These findings suggest that the exacerbation of cardiovascular pathology was not driven by the APOL1-G1 expression. As there was substantial atherosclerosis and calcification in all APOL1-transgenic mice at 9-months of age, differences among groups might have been present at time points that were not studied.

Importantly, this transgenic mouse model does not manifest atherosclerosis of coronary arteries as seen in humans. Few studies have successfully established coronary artery atherosclerosis in rodents. Mice are resistant to atherosclerosis and this may be partially due to higher HDL and lower LDL cholesterol levels.^75^ Although coronary plaque rupture has been difficult to model in mice,^75^ we confirmed APOL1 accumulation in atherosclerotic lesions, similar to that seen in humans.^21^ Further, none of the mice in the current study developed kidney disease, as assessed by normal urine albumin excretion and lack of renal pathology. Therefore, this mouse model is a suitable model to study cardiovascular disease with human APOL1 variant, uncomplicated by effects of chronic kidney disease. All of Us data supported these mouse findings, as APOL1-G1 variant was not associated with increased prevalence of atherosclerosis by multi-variate logistic regression analysis, adjusted for age, sex, and eGFR.

There are several limitations in this study. First, mice had mixed genetic backgrounds as described in the Methods. It might be the reason for the heterogeneity of phenotypes observed. Second, single-cell RNA-seq experiments include data from the two mice, one mouse from each genotype for comparison, due to resource limitations. This finding of hydrocephalus in these mice was incidental and warrants further investigation; additional human cohort data might strengthen this observation.

In conclusion, the central finding reported here is that the *APOL1-G1* risk variant is associated with hydrocephalus, possibly driven by mTORC2 activation. Better understanding of the prevalence and mechanisms of this condition may improve its management and outcomes.

## Acknowledgements

We thank the DNA Sequencing and Genomics Core (NHLBI/NIH) for sequencing support, Drs. Milton Pryor and Jingrong Tang (NHLBI/NIH) for experimental support, Dr. Zhenzhong Cui for mouse CSF collection procedure, Drs. Kris Ylaya, Joon-Yong Chung and Stephen Hewitt (NCI/NIH) for whole slide scanning, Dr. Luis Menezes for critical manuscript review. We appreciate receiving BAC/APOL1 mice from Merck & Co., Inc., Rahway, NJ, USA. We appreciate receiving APOL1 antibodies from Genentech, South San Francisco, CA. This work used the resources of the NIH HPC Biowulf cluster (http://hpc.nih.gov), the NIDDK Advanced Light Microscopy & Image Analysis Core (ALMIAC) and the NIDDK Mouse Transgenic Core Facility and NIH Mouse Imaging Facility (NINDS/NIH). We used BioRender.com to create figures. Part of this work was presented at the 16^th^ Meeting of the Hydrocephalus Society and the American Society of Nephrology Annual Meeting 2024.

The All of Us Research Program is supported by the National Institutes of Health, Office of the Director: Regional Medical Centers: 1 OT2 OD026549; 1 OT2 OD026554; 1 OT2 OD026557; 1OT2 OD026556; 1 OT2 OD026550; 1 OT2 OD 026552; 1 OT2 OD026553; 1 OT2 OD026548; 1OT2 OD026551; 1 OT2 OD026555; IAA #: AOD 16037; Federally Qualified Health Centers: HHSN 263201600085U; Data and Research Center: 5 U2C OD023196; Biobank: 1 U24OD023121; The Participant Center: U24 OD023176; Participant Technology Systems Center: 1U24 OD023163; Communications and Engagement: 3 OT2 OD023205; 3 OT2 OD023206; and Community Partners: 1 OT2 OD025277; 3 OT2 OD025315; 1 OT2 OD025337; 1 OT2 OD025276. In addition, the All of Us Research Program would not be possible without the partnership of its participants. The content of this publication does not necessarily reflect the views or policies of the Department of Health and Human Services, nor does mention of trade names, commercial products, or organizations imply endorsement by the U.S. Government.

## Declarations

### Ethics statement

Animal study proposal numbers are K097-KDB-17 & K096-KDB-20 (IACUC/NIDDK).

### Author Contributions

TY and JBK conceived the study design. TY and SS conducted mouse experiments. ZY, TY, VR, SS and AL dissected mouse aortas and quantified atherosclerosis. PB, DN and HW quantified aortic calcification. MFS and VJH conducted necropsy of mice. TY conducted single-nuclear RNA-seq capture and RNA-seq data analysis. SA and BB provided deidentified human brain tissue. DAS conducted echocardiogram. JM conducted MRI. ZXY assessed cardiovascular pathology. TY conducted the analysis of All of Us Cohort. TY drafted the manuscript, and all the authors edited the manuscript.

### Data Availability Statement

Original transcriptomic data have been deposited in GEO (GSE252599). Other data are available from the authors upon request.

