## Supplementary Material for "Apolipoprotein-L1 G1 variant contributes to hydrocephalus but not to atherosclerosis in apolipoprotein-E knock-out mice"

Running Title: Hydrocephalus in APOL1-G1/ApoE-KO mice


Teruhiko Yoshida[1,2], Zhi-Hong Yang[3], Shinji Ashida[4], Zu Xi Yu[3], Shashi Shrivastav[1], Krishna Vamsi Rojulpote[3], Piroz Bahar[3], David Nguyen[3], Danielle A. Springer[3], Jeeva Munasinghe[5], Matthew F. Starost[6], Victoria J. Hoffmann[6], Avi Z. Rosenberg[7], Bibi Bielekova[4], Han Wen[3], Alan T. Remaley[3], Jeffrey B. Kopp[1]

[1] National Institute of Diabetes and Digestive and Kidney Diseases, NIH, Bethesda, MD,

[2] Graduate School of Medicine, The University of Tokyo, Tokyo, JAPAN,

[3] National Heart, Lung, and Blood Institute, NIH, Bethesda, MD,

[4] National Institute of Allergy and Infectious Diseases, NIH, Bethesda, MD,

[5] National Institute of Neurological Disorders and Stroke, NIH, Bethesda, MD,

[6] Office of the Director, NIH, Bethesda, MD

[7] Department of Pathology, Johns Hopkins Medical Institutions, Baltimore, MD

*Corresponding author and lead contact:* Teruhiko Yoshida, MD PhD;

*address:* 7-3-1 Hongo, Bunkyo-ku, Tokyo, JAPAN 113-8655

**Table of Contents**

**Supplemental Tables**

Supplemental Table 1. Metrics summary of single-nuclear RNA-seq

Supplemental Table 2. Concept identifiers of hydrocephalus used in All of Us analysis

Supplemental Table 3. Concept identifiers of atherosclerosis used in All of Us analysis

Supplemental Table 4. Key resource tables

**Supplemental Figures**

Supplemental Figure 1. Echocardiography images.

Supplemental Figure 2. Downstream factors of RICTOR showing differential expression and predicted activation of RICTOR from single-nuclear RNA-seq data from choroid plexus epithelial cells, comparing APOL1-G1 mice and WT mice.

Supplemental Figure 3. CSF cytokine panel results

Supplemental Figure 4. Weight and albuminuria levels of BAC/APOL1xApoE-KO mice among APOL1 genotypes

Supplemental Figure 5. Hydrocephalus prevalence among APOL1 genotypes in the All of Us population cohort.

Supplemental Figure 6. Atherosclerosis prevalence among APOL1 genotypes in the All of Us population cohort.

Supplemental Figure 7. BAC/APOL1-G1 genome insertion site and neighboring gene expressions in single-nucleus RNA-seq data.

### Supplemental Table 1. Metrics summary of single-nuclear RNA-seq

| *Sample* | *Estimated Number of Cells* | *Mean Reads per Cell* | *Median Genes per Cell* | *Number of Reads* | *Valid Barcodes* | *Sequencing Saturation* | *Q30 Bases in Barcode* | *Q30 Bases in RNA Read* | *Q30 Bases in UMI* | *Reads Mapped to Genome* | *Reads Mapped Confidently to Genome* | *Reads Mapped Confidently to Intergenic Regions* | *Reads Mapped Confidently to Intronic Regions* | *Reads Mapped Confidently to Exonic Regions* | *Reads Mapped Confidently to Transcriptome* | *Reads Mapped Antisense to Gene* | *Fraction Reads in Cells* | *Total Genes Detected* | *Median UMI Counts per Cell* |
|---|---|---|---|---|---|---|---|---|---|---|---|---|---|---|---|---|---|---|---|
| 1_WT | 6,636 | 104,498 | 1,695 | 693,447,828 | 94.80% | 80.20% | 96.80% | 94.50% | 96.50% | 93.50% | 90.40% | 6.50% | 45.80% | 38.20% | 45.20% | 38.60% | 47.70% | 26,544 | 2,977 |
| 2_G1 | 15,626 | 83,902 | 1,425 | 1,311,056,762 | 95.00% | 84.20% | 96.70% | 94.20% | 96.50% | 94.00% | 90.90% | 6.70% | 42.90% | 41.40% | 46.40% | 37.70% | 51.40% | 26,313 | 2,303 |

**Supplemental Table 2. Concept identifiers of hydrocephalus used in All of Us analysis**

| Concept name (Condition) | Concept identifier |
| --- | --- |
| Acquired hydrocephalus | 4256924 |
| Benign intracranial hypertension | 312902 |
| Communicating hydrocephalus | 440700 |
| Congenital hydrocephalus | 438244 |
| Fetal hydrocephalus | 37204822 |
| Hydrocephalus | 4043738 |
| Non-obstructive hydrocephalus | 4105343 |
| Normal pressure hydrocephalus | 432899 |
| Obstructive hydrocephalus | 440385 |

**Supplemental Table 3. Concept identifiers of atherosclerosis used in All of Us analysis**

| Concept name (Condition) | Concept identifier |
|---|---|
| Atherosclerosis | 44825446 |
| Atherosclerosis | 1569271 |
| Atherosclerosis of aorta | 44823124 |
| Atherosclerosis of aorta | 35207841 |
| Atherosclerosis of coronary artery bypass graft(s) without angina pectoris | 45567167 |
| Atherosclerosis of coronary artery without angina pectoris | 764123 |
| Atherosclerosis of native arteries of extremities with intermittent claudication | 1569274 |
| Atherosclerosis of native arteries of the extremities | 44828992 |
| Atherosclerosis of native arteries of the extremities | 1569272 |
| Atherosclerosis of native arteries of the extremities with intermittent claudication | 44825447 |
| Atherosclerosis of other coronary vessels without angina pectoris | 1569146 |
| Coronary arteriosclerosis | 317576 |
| Coronary atherosclerosis | 44835931 |
| Coronary atherosclerosis due to calcified coronary lesion | 45567168 |
| Coronary atherosclerosis of native coronary artery | 44830080 |
| Coronary atherosclerosis of unspecified type of vessel, native or graft | 44835932 |
| Other and unspecified atherosclerosis | 1569320 |
| Unspecified atherosclerosis | 45581810 |
| Unspecified atherosclerosis of native arteries of extremities | 1569273 |

#### Supplemental Table 4. Key resource tables

| REAGENT or RESOURCE | SOURCE | IDENTIFIER |
|---|---|---|
| **Antibodies** | | |
| APOL1 | Genentech | 5.17D12, 3.7D6+3.1C1[1] |
| pNKCC1 | Millipore-Sigma | ABS1004 |
| pSPAK | Millipore-Sigma | 07-2273 |
| Phospho-SGK1 (Thr256) | Thermo Fisher Scientific | 44-1260G |
| pAkt | Cell Signaling | 9271 |
| Goat anti-Rabbit IgG (H+L) Highly Cross-Adsorbed Secondary Antibody, Alexa Fluor Plus 488 | Thermo Fisher Scientific | A32731TR |
| SGK1 | Abcam | ab43606 |
| β-actin | Santa Cruz Biotechnology | 47778 |
| **Chemicals, peptides, and recombinant proteins** | | |
| Doxycycline hydrochloride | Millipore-Sigma | D3072 |
| **Critical commercial assays** | | |
| Cholesterol E kit | Fujifilm Wako Chemicals | NC9138103 |
| ImmPRESS HRP Horse Anti- rabbit IgG Polymer Kit, Peroxidase | Vector Laboratories | MP-7401 |
| ImmPACT DAB EqV Peroxidase (HRP) Substrate | Vector Laboratories | SK-4103 |
| Lipofectamine 3000 | Thermo Fisher Scientific | L3000001 |
| Mouse Cytokine/Chemokine 32-Plex Discovery Assay Array | Eve technologies | MD32 |
| Chromium Next GEM Single Cell 3' Reagent Kit, v3.1 chemistry | 10xGenomics | 1000128 |
| **Deposited data** | | |
| Raw and analyzed single-nuclear RNA-seq data | This paper | GEO: GSE252599 |
| **Experimental models: Cell lines** | | |
| Human choroid plexus epithelial cells | ScienCell | 1310 |
| **Experimental models: Organisms/strains** | | |
| BAC/APOL1-G0, G1 | MSD<br>Okamoto et al.[2], Ryu et al.[3], Wakashin et al.[4] | N/A |
| ApoE-KO (B6.129P2-Apoe$^{tm1Unc}$/J) | Jackson Laboratory | 002052 |
| **Recombinant DNA** | | |
| TRE-Empty, TRE-APOL1-G0, G1, G2 plasmids | This paper | N/A |
| pCMV-Tet3G plasmid | Takara Bio USA | 631335 |
| **Software and algorithms** | | |
| CellRanger | Zheng et al.[5] | https://www.10xgenomics.com/support/software/cell-ranger |
| SoupX | Young et al.[6] | https://github.com/constantAmateur/SoupX |

| | | |
|---|---|---|
| DoubletFinder | McGinnis et al.[7] | https://github.com/chris-mcginnis-ucsf/DoubletFinder |
| Seurat | Hao et al.[8] | https://satijalab.org/seurat/ |
| SeuratData | Hao et al.[8] | https://satijalab.org/seurat/ |
| Ingenuity Pathway Analysis (IPA) | QIAGEN<br>Kramer et al.[9] | N/A |

#### Supplementary Figure 1

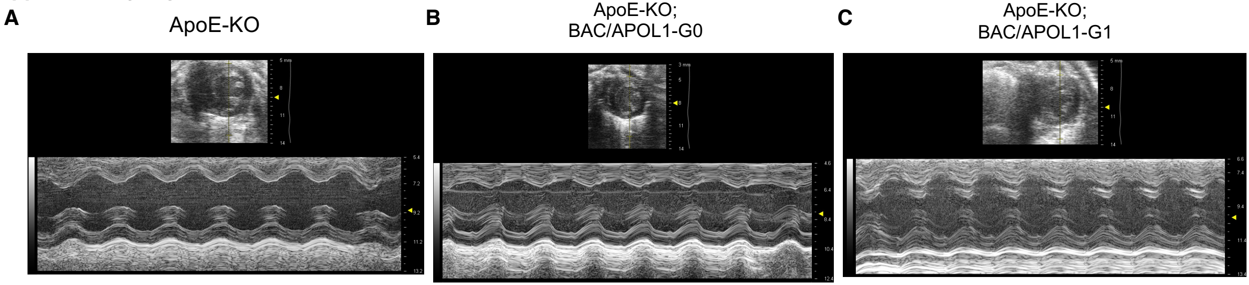


**Supplemental Figure 1. Echocardiography images**
(**A-C**) Representative M-mode images of echocardiography showed no difference among mice of each genotype.

#### Supplementary Figure 2

**A**

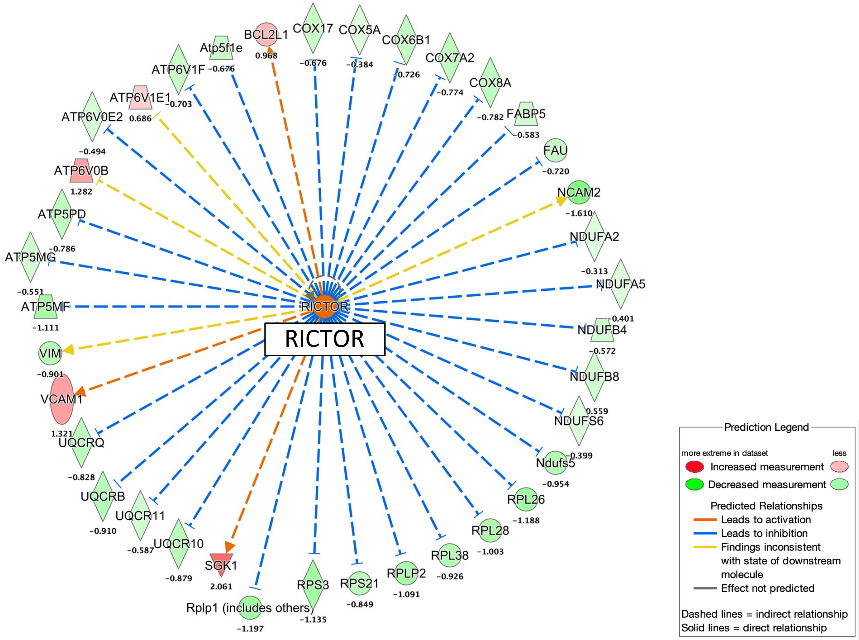


**Supplemental Figure 2. Downstream factors of RICTOR showing differential expression and predicted activation of RICTOR from single-nuclear RNA-seq data from choroid plexus epithelial cells, comparing ApoE-KO; BAC/APOL1-G1 mice and ApoE-KO mice.**

(**A**) Downstream factors of RICTOR indicated activation of RICTOR in choroid plexus epithelial cells of ApoE-KO; BAC/APOL1-G1 mice compared with ApoE-KO mice.

Number denotes the expression in log2 fold-change comparing ApoE-KO; BAC/APOL1-G1 mice and ApoE-KO mice.

Red denotes upregulation in ApoE-KO; BAC/ APOL1-G1, and green denotes downregulation in ApoE-KO; BAC/ APOL1-G1 mice.

**Supplementary Figure 3**

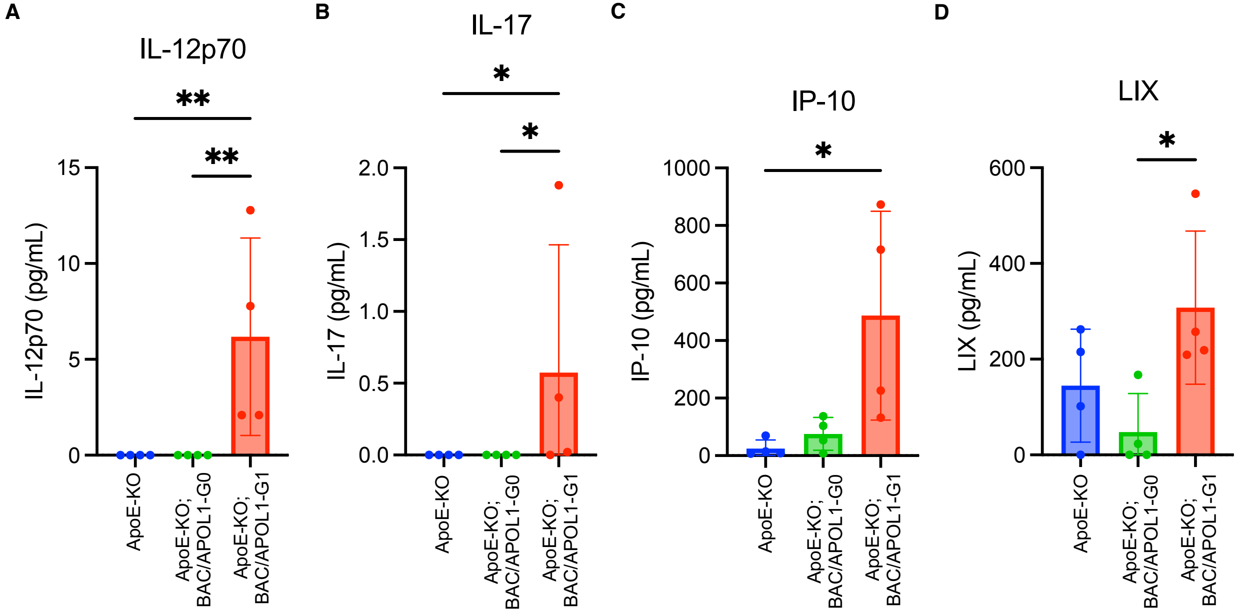


**Supplemental Figure 3. CSF cytokine panel results**
(**A-D**) Cytokines (IL-12p70, IL-17, IP-10 and LIX) measured in CSF showed higher levels in APOL1-G1 mice.
IP-10, Interferon gamma-induced protein 10 (CXCL10); LIX, LPS-induced CXC chemokine (CXCL5)

**Supplementary Figure 4**

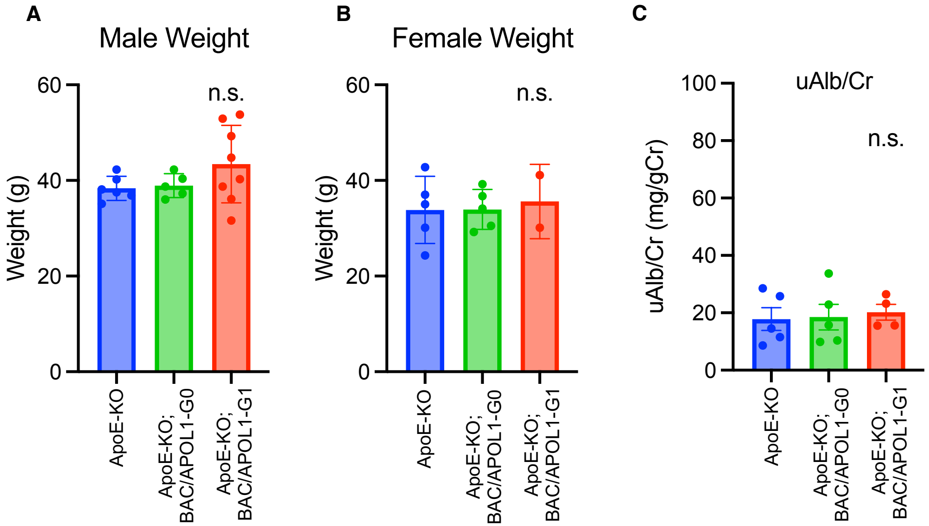


**Supplemental Figure 4. Weight and albuminuria levels of BAC/APOL1xApoE-KO mice among APOL1 genotypes**
(**A-B**) Weights of 9 month-old mice showed no difference among genotypes.
(**C**) Urinary albumin/creatinine ratio showed no difference among genotypes.

**Supplemental Figure 5**

**A**

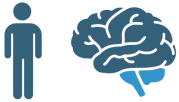

| | | Subjects | Case | Prevalence | Odds Ratio [95% CI] | P-value |
|---|---|---|---|---|---|---|
| Models | G1 allele recessive | | | | **1.73 [1.11-2.60]** | **0.012*** |
| | G1 allele dominant | | | | **1.30 [1.03-1.64]** | **0.025*** |
| | Risk allele recessive | | | | 1.19 [0.85-1.64] | 0.27 |
| | Risk allele dominant | | | | 1.19 [0.93-1.52] | 0.16 |
| APOL1 Genotype | G1/G1 | 2844 | 26 | 0.91% | **1.85 [1.16-2.86]** | **0.0092**** |
| | G1 or G2/G2 | 4505 | 21 | 0.47% | 0.94 [0.56-1.51] | 0.91 |
| | G0/G2 | 9594 | 43 | 0.45% | 0.90 [0.62-1.30] | 0.60 |
| | G0/G1 | 15382 | 100 | 0.65% | 1.31 [0.99-1.74] | 0.057 |
| | G0/G0 | 22172 | 110 | 0.50% | Ref | Ref |

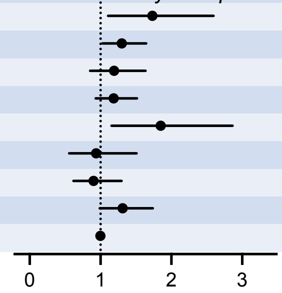


**B**

| Univariate analysis | *All (n=54497)* | *Hydrocephalus (n=300)* | *No hydrocephalus (n=54197)* | *P-value* |
|---|---|---|---|---|
| Age (mean, SD) | 53.0, 14.8 | 49.8, 14.4 | 53.0, 14.8 | 0.00021* |
| Male (n, %) | 23292, 43% | 44, 15% | 23252, 43% | 2.2E-16* |
| eGFR (mean, SD) | 75.5, 30.0 | 63.2, 25.9 | 75.6, 30.0 | 0.0048* |
| G1 allele recessive (n, %) | 2844, 5.2% | 26, 8.7% | 2818, 5.2% | 0.012* |
| G1 allele dominant (n, %) | 21590, 40% | 138, 46% | 21456, 40% | 0.025* |

**C**

<u>APOL1-G1 recessive model</u>

| Multi-variate logistic regression analysis | *Coefficient* | *Odds ratio [95% CI]* | *P-value* |
|---|---|---|---|
| Age (mean, SD) | -0.030 | 0.97 [0.95-0.99] | 0.0034* |
| Male (n, %) | -0.75 | 0.48 [0.21-0.95] | 0.048* |
| eGFR (mean, SD) | -0.014 | 0.99 [0.98-0.99] | 0.0032* |
| G1 allele recessive (n, %) | 1.471 | 4.35 [1.93-8.88] | 0.00014* |

**D**

<u>APOL1-G1 dominant model</u>

| Multi-variate logistic regression analysis | *Coefficient* | *Odds ratio [95% CI]* | *P-value* |
|---|---|---|---|
| Age (mean, SD) | -0.032 | 0.97 [0.95-0.99] | 0.0017* |
| Male (n, %) | -0.74 | 0.48 [0.22-0.96] | 0.0498* |
| eGFR (mean, SD) | -0.015 | 0.99 [0.98-0.99] | 0.0026* |
| G1 allele dominant (n, %) | 1.35 | 3.86 [2.09-7.50] | 0.000028* |

**Supplemental Figure 5. Hydrocephalus prevalence among APOL1 genotypes in the All of Us population cohort.**
(**A, B**) The prevalence of hydrocephalus stratified by APOL1 genotype and by genetic models and univariate analyses were shown. Fisher's exact test was used for APOL1 models, genotypes, and sex. t-test was used for age and eGFR.
(**C, D**) Multi-variate logistic regression analysis results for hydrocephalus odds ratios, adjusted for age, sex, and eGFR were shown with APOL1-G1 recessive model, and dominant model.

**Supplemental Figure 6**

**A**

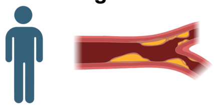

| | | *Subjects* | *Case* | *Prevalence* | *Odds Ratio [95% CI]* | *P-value* |
|---|---|---|---|---|---|---|
| Models | G1 allele recessive | | | | **1.19 [1.02-1.37]** | **0.024*** |
| | G1 allele dominant | | | | 0.98 [0.92-1.06] | 0.65 |
| | Risk allele recessive | | | | 1.04 [0.94-1.15] | 0.46 |
| | Risk allele dominant | | | | 0.99 [0.93-1.07] | 0.83 |
| APOL1 Genotype | G1/G1 | 2844 | 210 | 7.38% | **1.17 [1.00-1.36]** | **0.048**** |
| | G2/G2 | 1141 | 60 | 5.26% | 0.81 [0.61-1.06] | 0.13 |
| | G1/G2 | 3364 | 212 | 6.30% | 0.99 [0.85-1.15] | 0.88 |
| | G0/G2 | 9594 | 631 | 6.58% | 1.03 [0.93-1.14] | 0.53 |
| | G0/G1 | 15382 | 938 | 6.10% | 0.95 [0.87-1.04] | 0.25 |
| | G0/G0 | 22172 | 1417 | 6.39% | Ref | Ref |

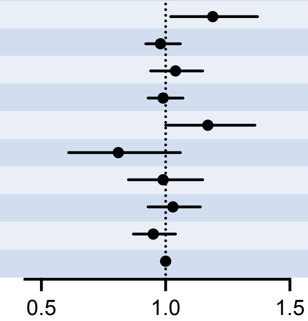


**B**

| Univariate analysis | *All (n=54497)* | *Atherosclerosis (n=3369)* | *No atherosclerosis (n=51128)* | *P-value* |
|---|---|---|---|---|
| Age (mean, SD) | 53.0, 14.8 | 65.4, 10.7 | 52.1, 14.6 | <2.0E-16* |
| Male (n, %) | 23292, 43% | 1512, 45% | 21784, 43% | 0.3 |
| eGFR (mean, SD) | 75.5, 30.0 | 69.0, 31.1 | 76.9, 29.6 | 9.7E-9* |
| G1 allele recessive (n, %) | 2844, 5.2% | 210, 6.2% | 2634, 5.2% | 0.024* |
| G1 allele dominant (n, %) | 21590, 40% | 1361, 40% | 20233, 40% | 0.65 |

**C**

#### APOL1-G1 recessive model

| Multi-variate logistic regression analysis | *Coefficient* | *Odds ratio [95% CI]* | *P-value* |
|---|---|---|---|
| Age (mean, SD) | 0.078 | 1.08 [1.07-1.09] | 5.5E-63* |
| Male (n, %) | 0.40 | 1.5 [1.22-1.83] | 8.6E-05* |
| eGFR (mean, SD) | -0.0072 | 0.993 [0.990-0.996] | 2.6E-05* |
| G1 allele recessive (n, %) | 0.21 | 1.23 [0.78-1.88] | 0.36 |

**D**

#### APOL1-G1 dominant model

| Multi-variate logistic regression analysis | *Coefficient* | *Odds ratio [95% CI]* | *P-value* |
|---|---|---|---|
| Age (mean, SD) | 0.078 | 1.08 [1.07-1.09] | 5.3E-63* |
| Male (n, %) | 0.41 | 1.50 [1.23-1.83] | 7.8E-05* |
| eGFR (mean, SD) | -0.0071 | 0.993 [0.990-0.996] | 2.7E-05* |
| G1 allele dominant (n, %) | 0.14 | 1.15 [0.95-1.40] | 0.16 |

**Supplemental Figure 6. Atherosclerosis prevalence among APOL1 genotypes in the All of Us population cohort.**
(**A, B**) The prevalence of atherosclerosis stratified by APOL1 genotype and by genetic models and univariate analyses were shown. Fisher's exact test was used for APOL1 models, genotypes, and sex. t-test was used for age and eGFR.
(**C, D**) Multi-variate logistic regression analysis results for atherosclerosis odds ratios, adjusted for age, sex, and eGFR were shown with APOL1-G1 recessive model, and dominant model.

**Supplemental Figure 7**

**A**

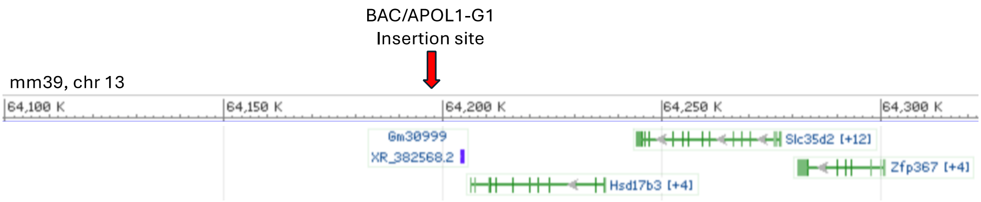


**B**

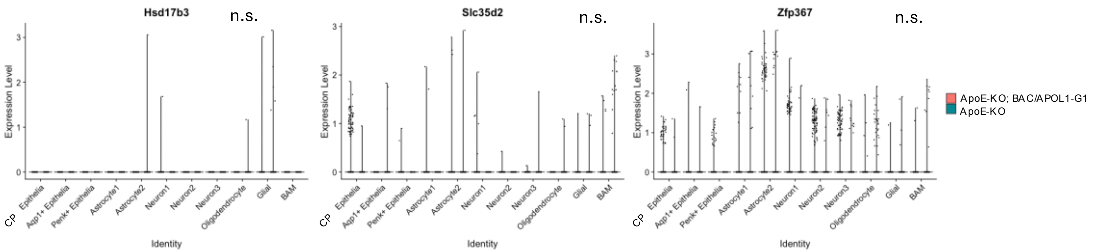


**Supplemental Figure 7. BAC/APOL1-G1 genome insertion site and neighboring gene expressions in single-nucleus RNA-seq data.**
(**A**) BAC/APOL1-G1 transgene insertion site was detected (mm39, chr13: 64,197,419) as shown.
(**B**) Volcano plots showed gene expression of neighboring genes (*Hsd17b3, Slc35d2, Zfp367*) around the transgene insertion site in choroid plexus epithelial cells from single-nucleus RNA-seq data.
